# Non-native crustacean predators conditionally trigger inducible defense behaviors in blue mussels

**DOI:** 10.64898/2025.12.04.691823

**Authors:** Ruxin Dai, Christopher D. Wells

**Affiliations:** Schiller Coastal Studies Center, Bowdoin College, Harpswell, ME, USA; Biology Department, Bowdoin College, Brunswick, ME, USA; Mathematics Department, Bowdoin College, Brunswick, ME, USA

## Abstract

Climate change is driving the poleward expansion of many species, including the Atlantic blue crab (*Callinectes sapidus*), whose northern range now extends into the Gulf of Maine. As a recently arrived predator, the blue crab may pose a threat to native prey communities already impacted by the earlier introduction of the European green crab (*Carcinus maenas*). This study investigates the defensive capacity of native blue mussels (*Mytilus edulis*) in response to 45 h of exposure to chemical cues of blue versus green crabs. Mussels were hypothesized to invest more energy in defensive behaviors in response to green crab cues, owing to a longer shared ecological history in the Gulf of Maine. The results support this hypothesis: female blue mussels exposed to green crab cues spawned, and small mussels exhibited increased relocation behavior. In contrast, exposure to blue crab cues did not alter mussel reproductive, movement, or attachment behaviors. Notably, neither treatment significantly affected mussel attachment as measured by byssal thread quantity and tenacity, suggesting that this behavior may not be a primary defense mechanism against crab predators in this context, contrary to previous findings. These findings underscore the importance of understanding species- and size-specific interactions as expanding ranges and species introductions continue to reshape predator-prey dynamics across coastal ecosystems.

## Introduction

Climate change driven increases in sea surface temperatures are reshaping coastal ecosystems worldwide (Walther et al. 2002, Harley et al. 2006). As waters warm, many species have extended their ranges poleward, disrupting established trophic structures in regions outside their historical distributions (Parmesan & Yohe 2003). However, the rate and direction of these shifts vary among taxa in response to local temperature changes (Pinsky et al. 2013).

One of the most pronounced examples of such ecological change is occurring in the Gulf of Maine, which ranks among the fastest-warming marine systems on Earth (Pershing et al. 2021). This environmental shift has enabled commercially valuable species, such as black sea bass (*Centropristis striata*), silver hake (*Merluccius bilinearis*), and Atlantic blue crabs (*Callinectes sapidus*), to expand their historic northern limits into this region (Nye et al. 2011, Johnson 2015, McBride et al. 2018). Although the presence of these species is likely to generate commercial revenue, the ecological consequences for Gulf of Maine systems remain uncertain.

The Atlantic blue crab (*C. sapidus*, herein referred to as blue crab) is of particular concern, given the precedent set by prior crustacean invasions. In the early 1800s, the European green crab (*Carcinus maenas*, herein referred to as green crab) was introduced to the U.S. Mid-Atlantic, and subsequently reached the Gulf of Maine by the early 1900s (Scattergood 1952). In their native European range, green crabs primarily consume bivalves but also feed on gastropods, annelids, and other crustaceans (Le Roux et al. 1990). In the Gulf of Maine, an ecosystem naïve to this predator, their foraging led to heavy reductions in the number of soft-shell clams (*Mya arenaria*) (Glude 1955, Tan & Beal 2015), and contributed to the rapid decline of eelgrass (*Zostera marina*) meadows (Matheson et al. 2016). Moreover, green crabs’ preference for rough periwinkle (*Littorina saxatilis*) over common periwinkle (*Littorina littorea*) has altered the intertidal distribution of these ecologically important gastropods (Lubchenco 1978, Eastwood et al. 2007).

Although blue and green crabs occupy different ecological niches and history of expansion into the Gulf of Maine, the green crab’s impact provides a useful analogue for anticipating the consequences of blue crab expansion in the Gulf of Maine (deRivera et al. 2005). Both green and blue crabs are opportunistic omnivores with a strong preference for bivalve prey (Ropes 1968, Seitz et al. 2011). This overlap raises concerns that a rising blue crab population could further pressure the already diminished bivalve communities. Additionally, blue crabs consume juvenile and molting American lobsters (*Homarus americanus*), the basis of the United States’ most economically valuable single-species fishery (Steinback et al. 2008, Meyer-Rust et al. 2024). The prospect of direct predation on lobster stock underscores both ecological and commercial implications of blue crab range expansion.

Importantly, blue crabs are larger, more aggressive, and more energetically demanding predators than their green crab counterparts (Millikin & Williams 1984), and engage in size-dependent agonistic and trophic interactions with them (deRivera et al. 2005, Smith et al. 2025). Although blue crabs are a climate-induced range-expanding species and green crabs are an invasive species, both types of novel predators can exert similarly disruptive pressures on recipient ecosystems (Sorte et al. 2010, Bellard et al. 2016). Consequently, understanding how local preys respond to the encounter with a range-expanding predator may yield insights into the broader dynamics of ecological novelty. Among potential prey species, blue mussels (*Mytilus edulis*, herein referred to as blue mussels), a shared prey of the two predators and dominant bivalve in the Gulf of Maine, offer a compelling model for investigating how a novel predator influences prey behavior and morphology.

Blue mussels serve as ecosystem engineers in temperate rocky intertidal communities, historically structuring much of the Gulf of Maine shoreline (Menge 1976). Their capacity to detect and respond to predator-derived chemical cues (i.e., kairomones) is well documented (Petraitis 1987, Smith & Jennings 2000, Lowen et al. 2013). Detection of threat triggers a series of induced defenses, ranging from immediate behavioral responses such as shell closure (Dzierżyńska-Białończyk et al. 2019), aggregation in clusters (Côté & Jelnikar 1999), and increased byssal thread production (Côté 1995) to long term physiological changes such as strengthening of adductor muscles (Freeman et al. 2009) and production of thicker shells (Sherker et al. 2017). These inducible defenses make blue mussels an ideal system for assessing how prey differentiate between familiar and novel predators under changing environmental conditions.

Of particular interest is byssal thread production (i.e., byssogenesis), which has been characterized as an anti-predator mechanism (Reimer & Tedengren 1996, Leonard et al. 1999, Cheung et al. 2004, Garner & Litvaitis 2013a). Byssal threads are extracellular collagenous fibers secreted by a specialized gland, anchoring mussels to the substratum. The quantity and quality of byssal threads respond nonlinearly to multiple environmental factors, including temperature, wave exposure, and predator presence (Hatcher et al. 1997, Garner & Litvaitis 2013b, Rickaby & Sinclair 2018). Moreover, predator-specific prey defenses are often shaped by the spatial and temporal history of interaction (Hemmi & Merkle 2009, Crawford et al. 2012). For example, mussels from populations naïve to green crabs did not increase byssal thread production upon exposure to green crab kairomones, whereas previously exposed populations did (Reimer & Harms-Ringdahl 2001, Rickaby & Sinclair 2018). This pattern reflects the broader naïve prey hypothesis, which posits that prey naïve to a given predator invests less energy in induced defenses, or fails to exhibit them altogether (Sih et al. 2010, Carthey & Banks 2014, Carthey & Blumstein 2018). Given the recent northward expansion of blue crabs, Gulf of Maine blue mussels may display varying degrees of naiveté toward this novel predator.

Naiveté could manifest as reduced energy allocation in byssogenesis. Because byssal threads are costly to produce, energy diverted toward their production cannot be used for somatic growth or reproduction (Roberts et al. 2021, Roberts & Carrington 2023). Alternatively, blue mussels can exhibit predator-induced movement, abandoning one location to seek refuge in another. Each relocation compounds energetic costs, as mussels must abandon existing byssal threads and synthesize new ones (Reimer & Tedengren 1997, Ishida & Iwasaki 2003). This cycle of detachment and reattachment represents a substantial energy expenditure for defense. Beyond byssogenesis and movement, mussels may also exhibit a distinct behavioral response under acute threat: induced gamete release, also known as “escape by gametes” (sensu Reimer, 1999). This behavior can occur when mussels spawn in response to a sudden stressor. In aquaculture, for example, heat shock is often used to induce spawning (Saurel et al. 2022). Rather than representing a strategic reallocation of energy, such spawning reflects a terminal investment strategy: an attempt to propagate genes before imminent mortality. Consequently, the resulting gametes often experience suboptimal survival conditions (Hennebicq et al. 2013, Mredul et al. 2024) and may be underdeveloped.

Since antipredator responses in *M. edulis* carry considerable energetic trade-offs, the strength or presence of these responses likely depends on the population’s history of exposure to a given predator. Because neither blue nor green crabs are native to the Gulf of Maine but differ markedly in their duration of co-occurrence with blue mussels – approximately 120 years for green crabs versus only a few years for blue crabs – it is plausible that mussels respond differently to kairomones from each predator. To test the role of co-occurrence history in mediating prey responses to invasive versus range-expanding predators, we compared the behavior of blue mussels from Midcoast Maine when exposed to blue crab and green crab kairomones. We used byssal thread production, relocation behavior, and reproductive output as proxies for energy investment in antipredator defenses. Consistent with the naïve prey hypothesis, we predicted that green crab cues would elicit a pronounced defensive response, whereas blue crab cues, being evolutionarily novel, would not.

## Materials and Methods

### Sample collection & husbandry

Live blue mussels were collected from a culvert downstream of a mudflat community at Strawberry Creek, Harpswell, ME, USA (43.8137 °N, 69.9444 °W). The site has had no reported sightings of blue crabs and none were observed during collection. Green crabs were present at this location. After collection, all epizoic growth was removed from shell exteriors, and mussels were left to acclimate for at least three days in laboratory conditions. To ensure that mussels with damaged byssal producing organs were excluded from the experiment, only mussels that have laid down threads in the 3-day acclimation period were kept and used in subsequent experiments. Mussels were collectively housed in the same flow through container (Coleman Performance 70-quart, 74 × 39 × 37 cm) with continuous 25-µm filtered seawater input. Mussels were only used once to ensure naïveté to blue crab scents. For the whole experimental period (October 14 to November 18, 2024), mussels were fed 4 times per week with 2-mL doses of a mixed-algae liquid shellfish feed (Reed Mariculture Shellfish Diet 1800). The coarse filtration (250 µm) of incoming seawater likely allowed mussels to access phytoplankton as an additional food source. Mussels used for the first four trials were collected on October 11, 2024, and mussels used for the eight remaining trials were collected on October 27, 2024.

Live blue crabs were collected between October 10 and November 14, 2024, at New Meadows Lake, ME, USA (43.9306 °N, 69.8624 °W) with baited blue crab traps (60 × 50 × 50 cm; Ketcham Supply Co., New Bedford, MA). New Meadows Lake is a soft-bottom brackish wetland environment upstream of the tidally driven New Meadows River. Each crab was individually housed in flow through containers that shared the same water source (51 × 30 × 20 cm). Blue crabs were reused due to logistical constraints.

Live green crabs were collected from three subtidal populations in October 2024. Baited cylindrical Blanchard-style traps (90 × 40 × 40 cm) were deployed near Bowdoin College’s Schiller Coastal Studies Center (43.7926 °N, 69.9545 °W) and downstream of Strawberry Creek (43.8120 °N, 69.9457 °W) where the blue mussels were collected. Additional green crabs were collected as bycatch in the blue crab traps from New Meadows Lake. Green crabs were held collectively in a perforated container (76 × 43 × 46 cm) off the floating dock at the Schiller Coastal Studies Center and reused during the experimental period. Both species of crabs were fed *ad libitum* on a constant supply of blue mussels and frozen menhaden (*Brevoortia patronus*).

Blue crabs and mussels were housed in filtered sea water at ambient environmental temperatures, which ranged from 6.3 to 12.9 °C during the experiment.

### Experimental Design

To test the hypothesis that blue mussels would produce stronger or more byssal threads in the presence of blue and green crabs, mussels were conditioned in water containing crab chemical cues for 45 h. Upstream of mussels, filtered sea water at ambient environmental temperature flowed through 66 L coolers (Coleman Performance 70-quart, 74 × 39 × 37 cm) with one of three treatments: filtered sea water (control), green crab effluent, and blue crab effluent. The two crab treatments were created by housing live crabs in containers during the experimental period. The treated effluent was then distributed downstream to clear plastic containers through water manifolds (one per treatment) where mussels were housed individually in small plastic containers (24 × 16 × 11 cm).

For the blue crab effluent, two blue crabs (average 1.0 kg total mass, 135-205 mm carapace width) were housed in one cooler, separated by a metal mesh divider to reduce intraspecific aggression. For the green crab effluent, the number of green crabs used in each trial was different, but green crab biomass was matched closely with the blue crab biomass (average 1.1 kg total mass, 50-80 mm carapace width). Green grabs were not divided within the cooler. Due to logistical constraints, crabs that sustained injuries (i.e., injured chelae or missing limbs) were included in the experiment. None of the crabs were injured to the extent that their locomotion was compromised. In addition, none of the organisms were fed during the experiment to avoid confounding scents.

In total, we ran 12 trials with a total of 68 replicates per treatment. For the first four trials (October 14 to October 23, 2024), 15 mussels of any size (15-90 mm) were randomly assigned a treatment (n = 5 per treatment). However, initial analyses indicated that mussel size may have an impact on their ability to generate byssal threads. Therefore, for the subsequent eight trials (November 1 to November 18), 18 mussels were separated into 3 size ranges (0-30 mm, 31-60 mm, 61-90 mm) and evenly distributed across treatments (n = 6 per treatment).

The number of byssal threads produced was determined by counting the number of plaques that were attached. Prior to the trials, all existing byssal threads were cut from the mussels to differentiate thread production during the experiment from previous efforts with care to not damage the byssal thread producing organ. Byssal threads produced during the trial but no longer attached to the mussel were counted separately. The number of byssal thread clusters and the number of byssal threads within each cluster were also counted. Clusters were defined as spatially distinct groupings of byssal thread plaques.

The attachment force of individuals was determined by attaching each blue mussel to a calibrated Mxmoonfree ZMF-500N force detector and manually pulling the mussels off their substratum (Babarro et al. 2008). In some cases, attachment force could not be measured either because the attachment force was below the sensitivity threshold of the detector or because the mussel was attached at a corner, preventing an even, perpendicular pull. These individuals were excluded from force analyses.

### Statistical Analysis

All data analyses and visualizations were conducted in R version 4.4.3 (R Core Team 2025).

To look at the effects of effluent treatment on overall attachment force and per byssal thread force, we fit a negative binomial generalized linear mixed-effects model (GLMM) with effluent treatment, number of byssal threads, water temperature, and mussel length as fixed factors (R package glmmTMB 1.1.10, Brooks et al. 2017). We also included interaction terms for treatment, water temperature, and mussel length, while treating trial number as a discrete random factor. Mussels that did not attach during the trial, attached but moved without producing byssal threads in their new position, or attached to parts of their container that precluded attachment to the force detector were excluded in these analyses.

The effect of effluent treatment on the number of byssal threads produced was also assessed with a negative binomial GLMM. Effluent treatment, water temperature, and mussel length were treated as fixed factors and trial as a random effect. All interaction terms were included. We assessed the number of byssal threads in two ways: (1) the number attached to the mussel to the substratum at the end of the trial and (2) the total number produced throughout the trial, regardless of whether they were still attached to the mussel. Additionally, we assessed the number of clusters and the median number of byssal threads per cluster using the same approach.

For all negative binomial GLMMs, when the variance of trial was near zero (σ^2^ ≪0.001), we removed trial from the model and ran a negative binomial regression (GLM) instead (R package MASS 7.3-61, Venables & Ripley 2002).

Initial results did not differ when the first four trials were included in the analysis in comparison to when the eight trials were analyzed independently, despite slight differences in sizes of mussels sampled and sample size. Thus, all trials were included in the above analyses.

## Results

In total, 154 of the 204 mussels (75%) laid byssal threads during their trial and 141 of the mussels (69%) were attached at the end of their trial. Of the attached individuals, 75 individuals (53%) were attached in areas of the container where the attachment force could be quantified.

Crab effluent from neither species had an impact on the amount of byssal threads produced or tenacity of the threads after 45 h of exposure (GLM, χ^2^ ≤ 5.6, *p* ≥ 0.06). However, crab effluent did have an interactive effect with mussel length on the number of byssal thread clusters produced (Figure 1A, GLM, χ^2^ = 8.0, *p* = 0.02). In that case, green crab effluent significantly increased the number of clusters produced when mussels were small (< 3 cm).

**Figure 1.**
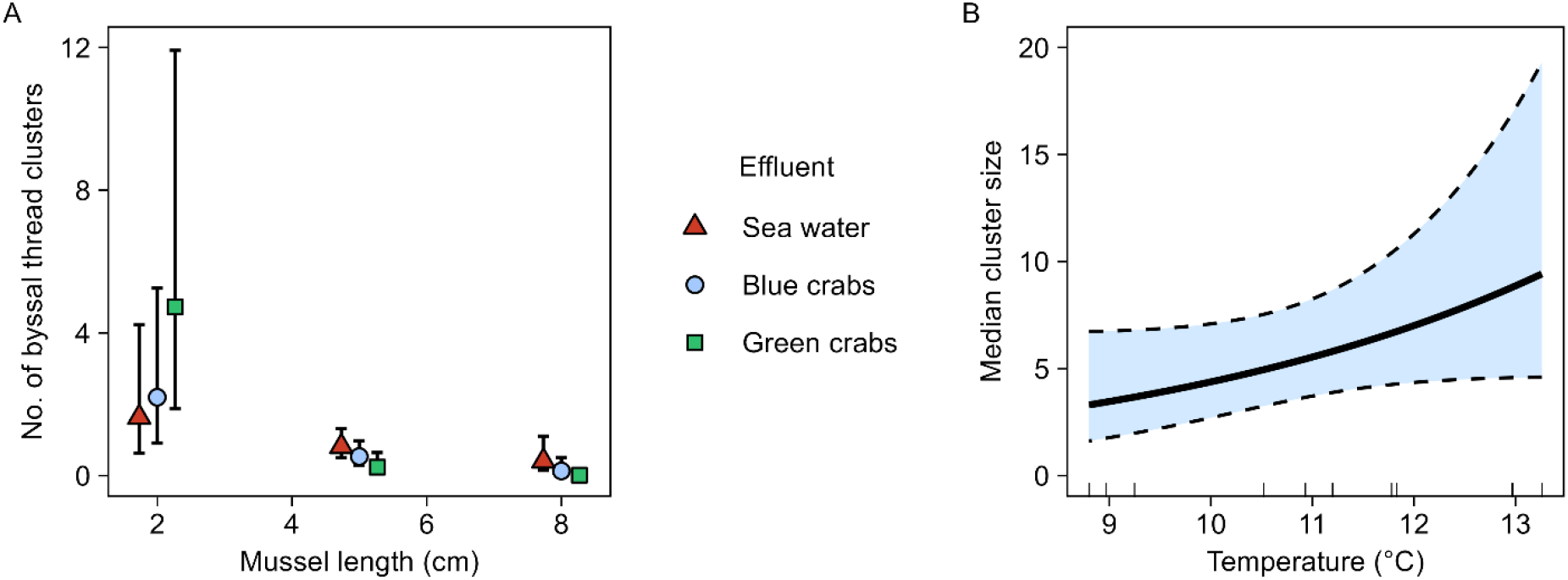
Number of byssal thread clusters (A) in response to mussel length and effluent treatment and (B) in response to changes in water temperature. (A) Error bars and (B) shaded regions bound by dashed lines indicate 95% confidence intervals. (A) Points are jittered around 2, 5, and 8 cm, so all points are visible. (B) The rug plot denotes observed data.

Interestingly, 11 of the 24 mussels (46%) exposed to green crab effluent in the last four trials (November 11 to 18, 2024) released yellow eggs (Figure S1 and S2 in the Supplement). Assuming an approximate 1:1 sex ratio in the collected mussels, this indicates that nearly all females in these trials released gametes. No mussels exposed to blue crabs or only seawater (control) spawned. It was unclear whether males also spawned. Mussel size did not seem to be a factor for spawning in our size range. Mussels that spawned ranged from 22 to 69 mm, the smallest to the largest size randomly assigned to green crab effluent during those trials. No mussels released eggs in the first eight trials (132 mussels; October 14 to November 8, 2024).

While crab effluent did not broadly influence byssogenesis, temperature and mussel size both affected production and attachment force of byssal threads. The number of attached byssal threads and the total production of byssal threads both increased with warmer water (Figure 2A and 2B, GLM, χ^2^ ≥ 13.2, *p* < 0.01) and total production decreased with mussel size (Figure 2C, GLM, χ^2^ = 11.3, *p* < 0.01). The number of attached byssal threads was not related to mussel length (GLM, χ^2^ = 1.8, *p* = 0.18). Median cluster size significantly increased with water temperature (Figure 1B, GLM, χ^2^ = 7.2, *p* = 0.01), but was not related to mussel length (GLM, χ^2^ = 0.5, *p* = 0.48). Total attachment force was higher when mussels were attached with more byssal threads and for larger mussels (Figure 3A and 3B, GLM, χ^2^ ≥ 29.4, *p* < 0.01), but was not related to water temperature (GLM, χ^2^ = 3.4, *p* = 0.07). The per byssal thread force was highest with larger mussels (Figure 3A, GLM, χ^2^ = 24.4, *p* < 0.01).

**Figure 2.**
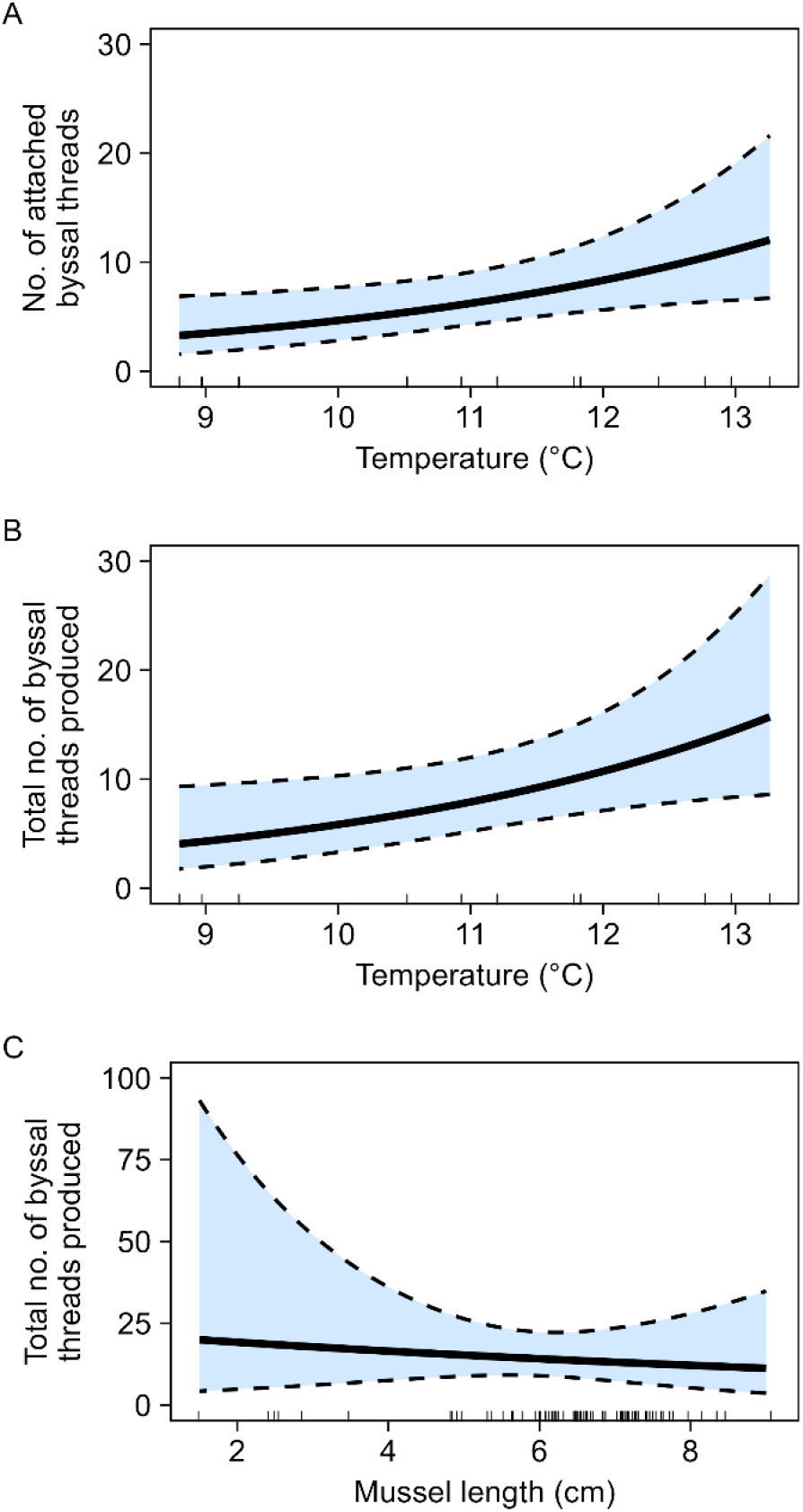
Number of (A) attached byssal threads and (B, C) total byssal threads produced in response to (A, B) temperature and (C) mussel length. The bold lines represent model predictions, while shaded regions bound by dashed lines indicate 95% confidence intervals. Rug plots denote observed data. The rug plot for mussel length was jittered so all measurements are visible.

**Figure 3.**
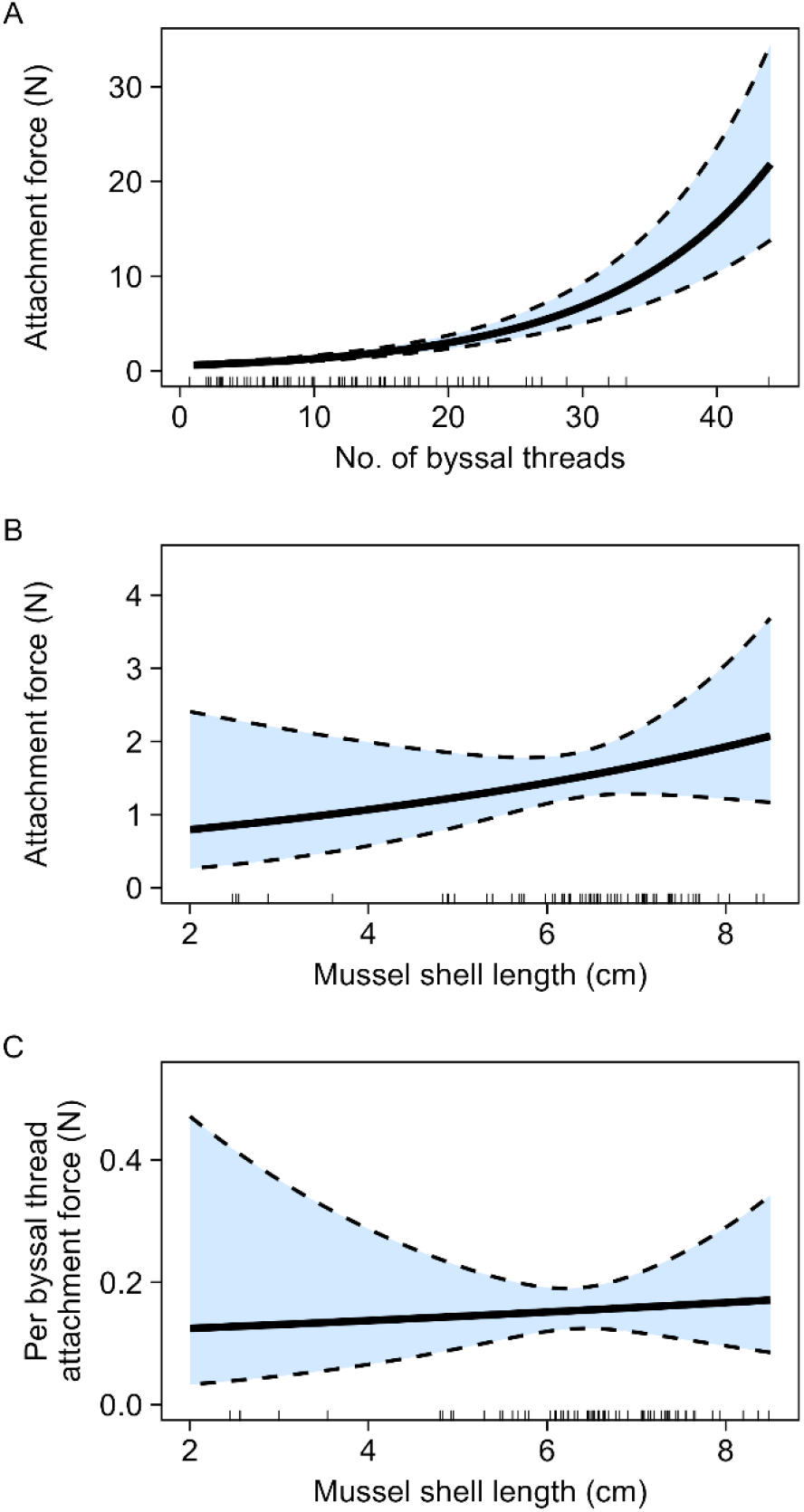
(A, B) Total and (C) per byssal thread attachment force in response to (A) number of byssal threads produced and (B, C) mussel shell length. The bold lines represent model predictions, while shaded regions bound by dashed lines indicate 95% confidence intervals. Rug plots denote observed data. Rug plots were jittered, so all measurements are visible.

## Discussion

Contrary to expectations, neither green nor blue crab effluent impacted blue mussel byssogenesis parameters after 45 hours of exposure. In contrast, green crab effluent affected behavior: small mussel movement increased (Figure 1A) and all observed spawning occurred in green crab treatments. Although past studies have demonstrated the ability of mussels to respond to predators by increasing thread quantity and tenacity, many were conducted under different environmental contexts. For example, they used different source populations (Reimer & Harms-Ringdahl 2001), different predators (Côté 1995, Reimer & Tedengren 1997), or were field experiments with uncontrolled covariates (Leonard et al. 1999). Predator-induced defenses in mollusks are neither taxonomically nor geographically ubiquitous. In this case, blue mussels in the Gulf of Maine are not responding to local blue crab or green crab populations by increasing their attachment to substrata but are changing their movement and reproductive behavior.

Variation in byssogenesis was instead modulated by abiotic and morphological factors. Byssal thread quantity increased with water temperature (Figure 1B, 2A, and 2B), a well-established pattern consistent with ectotherm bioenergetics (Young 1985). While greater thread production typically enhances attachment (Bell & Gosline 1996), especially since attachment force increases exponentially with thread quantity (Figure 3A), higher temperatures and thread production did not translate to stronger attachment in this study. Attachment depends on both thread quantity and quality. Here, individual thread tenacity showed no relationship with temperature, whereas in other work higher temperatures have reduced tenacity via malformed threads or faster denaturation (Lachance et al. 2008, Newcomb et al. 2019). Consequently, the net temperature effect on attachment is nonlinear, which has been found to vary across a 10–24 °C range. As oceans warm, temperature’s uneven effects on these parameters will complicate predictions of future mussel attachment strength.

Similarly, mussel size also had a non-linear relationship with attachment parameters. While the number of attached threads at 45 h was independent of mussel size, smaller mussels produced more total threads and more thread clusters, corroborating findings in *Mytilus galloprovincialis*. In contrast, larger mussels produced fewer clusters, indicating less frequent relocation and therefore reduced energy expenditure on movement. This difference in behavior may be driven by both physiological and ecological factors: larger mussels have higher basal metabolic rates, leaving less energy for movement (Sebens et al. 2018) and, above a threshold, grow into a size refuge from predation (Paine 1976, Seed 1980). With similarly sized clusters by the end of the experiment, larger mussels are more strongly attached because they can produce more tenacious threads (Figure 3C; Lee et al. 1990). Thus, mussel size influences how energy is allocated between movement, thread production, and thread quality.

On top of temperature and mussel size being constant modulators of byssogenesis, smaller mussels (∼2 cm) produced more thread clusters when exposed to green crab effluent (Figure 1A), which reflects both abandonment of prior attachment sites and relocation. Movement may be indicative of attempts to find conspecifics to form aggregations, a known anti-predator behavior in response to predator cues (Côté & Jelnikar 1999). Since this behavior involves energetically costly reinvestment in new byssal threads, increased movement in smaller mussels exposed to green crab cues suggest prioritization of exploring new attachment sites and finding conspecifics over strengthening attachment at a single site.

Additionally, exposure to green crab effluent coincided with all observed spawning events during the experiment and only during the final four trials (November 11-18). Phenology may be a factor: mussel gametes were likely underdeveloped during earlier trials. Blue mussels typically begin gonad redevelopment around October and November, continuing through the winter to full development by February (Brown 1967, Gosling 2003). The spawning activity elicited by green crabs during the early window of their reproductive cycle during these final trials may therefore represent a strategic energetic trade-off under perceived predation risk. Having already invested energy in gonadal development, individuals face a sunk cost, which favors completing spawning despite suboptimal timing rather than losing prior investment. This interpretation aligns with prior research showing that blue mussels exposed to predator cues increase their gonad-to-flesh ratio to maximize reproductive output (Reimer 1999). Conversely, spawning carries an opportunity cost, as the individual would have to reinvest in gonads or miss the optimal developmental window of the year.

Together, these observations suggest that the Gulf of Maine blue mussels can recognize and respond to green crab cues, particularly through increased movement and reproductive activity. Our results indicate that the Gulf of Maine blue mussel population has effectively “learned” to recognize green crab cues from an evolutionary perspective. Within the native range of both green crabs and blue mussels in the North Sea, blue mussels produce more byssal threads when exposed to green crab effluent (Reimer & Harms-Ringdahl 2001). However, in the adjacent, predator-free Baltic Sea, where crustacean predators have been absent for around 2000 years, mussels exhibited a weaker response (Bulnheim & Gosling 1988, Reimer & Harms-Ringdahl 2001). Locally, other mollusks also exhibit inducible defenses to green crab cues: dog whelks (*Nucella lapillus*) reduce foraging (Large et al. 2012) and the smooth periwinkle (*Littorina obtusata*) thickens its shell (Jacobsen & Stabell 1999). Our results join these examples in highlighting that prey responses are population specific and modulated by historic local predator exposure.

The lack of response to blue crab cues supports the naivete hypothesis which predicts that cue recognition depends on cohabitation history. Indeed, green crabs entered the Gulf of Maine long before blue crabs. Across animals, erosion of naiveté toward novel predators takes around 200 generations (Anton et al. 2020). We observed responses much sooner: with a generation time of one to two years, (Beyer et al. 2017) and roughly 120 years of coexistence with green crabs, the Gulf of Maine mussel population has evolved to respond in approximately 60-120 generations. This suggests rapid adaptation, and importantly, this response is not generalized across crab species.

Mussel responses therefore appear to be species-specific rather than triggered by a universal “crab cue.” In nearby locations in the Gulf of Maine, mussels do not elevate byssogenesis in the presence of rock crabs (*Cancer irroratus*) (Garner & Litvaitis 2013a). suggesting potential reliance on alternative defense strategies when confronted with crustacean predators. Thus, although the exact mechanisms behind these behaviors are yet unclear, changes in reproductive behavior and resource allocation may be alternative mechanisms. Differences in predation pressure, physiological constraints, trade-offs in energy allocation, and evolutionary history all likely play a role in observed variation in responses.

This study underscores how temperature and body size differentially affect movement frequency, thread number, and thread tenacity, and how predator cues can modulate behavior without altering standard byssogenesis metrics. Because spawning behavior can further modulate both byssal thread quantity and quality (Carrington 2002, Babarro & Reiriz 2010), multiple metrics are required to characterize attachment strategies. These findings contribute to growing evidence that predator-induced defenses in blue mussels are context-dependent rather than taxonomically or geographically universally consistent, complicating forecasts of population and ecosystem responses under climate change. If we cannot reliably predict how one key survival mechanism in a well-studied species will behave under warming conditions, anticipating broader ecosystem impacts becomes even more uncertain. Yet these uncertainties are crucial to confront.

Recognizing this complexity of behavioral responses underscores the importance of designing experiments that mirror natural conditions as closely as possible. The continuous flow of predator effluent used in this experiment likely approximated a more realistic cue exposure than previous laboratory studies that have relied on concentrated pulse addition of chemical cues. Observed sensitivity to green-crab cues in a flow-through system justifies follow-up studies in field or mesocosm settings with realistic hydrodynamics. Including tenacity, reproduction, and movement data, not thread count alone, yielded a fuller picture of how mussels balance risk, attachment, and energy allocation, with direct relevance to aquaculture, where attachment dynamics underpin rope-based culture (Roberts et al. 2021).

Looking forward, several avenues are poised for further investigation. Comparative studies across other economically and ecologically important bivalves, such as oysters, scallops, and clams, could help further understanding on whether similar forms of behavioral plasticity exist in other taxa under predator or environmental stress in their respective anti-predator strategies. Do these species exhibit analogous plasticity in attachment or alternative defenses? Moreover, future research could explore the genetic or molecular underpinnings of predator-specific byssogenesis, clarifying whether responses are conserved or species-specific. Long-term exposure experiments will also help determine if mussels shift strategies over time or become desensitized to predator cues. As climate change and range expansions reshape communities, understanding the variability and adaptability of attachment behaviors will be key to predicting ecosystem resilience and supporting both conservation and aquaculture goals.

## Acknowledgements

This work was made possible by Bowdoin College’s Schiller Coastal Studies Semester program and support of the staff at Schiller Coastal Studies Center: Heidi Franklin, Holly Parker, Jaret Reblin, Clinton Thompson, and Joe Tourtelotte. We thank the Schiller Coastal Studies Semester cohort, particularly Reuben Siegel and Layla Silva for coordinating transportation logistics and their patience in the animal husbandry process. We also extended our gratitude to Jessie Batchelder from Manomet Fisheries for her advice and company in the blue crab collection. Trapping was conducted under ME-DMR SL-2024-11-04 EDU. This research was made possible with support from Peter J. Grua and Mary G. O’Connell Student Research Fund and the Bowdoin College Biology Department Student Research Fund.

## References

Anton A, Geraldi NR, Ricciardi A, Dick JTA (2020) Global determinants of prey naiveté to exotic predators. Proceedings of the Royal Society B: Biological Sciences 287:20192978

Babarro JMF, Fernández Reiriz MJ, Labarta U (2008) Secretion of byssal threads and attachment strength of Mytilus galloprovincialis: The influence of size and food availability. Journal of the Marine Biological Association of the United Kingdom 88:783–791

Babarro JMF, Reiriz MJF (2010) Secretion of byssal threads in Mytilus galloprovincialis: Quantitative and qualitative values after spawning stress. Journal of Comparative Physiology B 180:95–104

Bell EC, Gosline JM (1996) Mechanical design of mussel byssus: Material yield enhances attachment strength. Journal of Experimental Biology 199:1005–1017

Bellard C, Cassey P, Blackburn TB (2016) Alien species as a driver of recent extinctions. Biology Letters 12:20150623

Beyer J, Green NW, Brooks S, Allan IJ and others (2017) Blue mussels (Mytilus edulis spp.) as sentinel organisms in coastal pollution monitoring: A review. Marine Environmental Research 130:338–365

Brooks ME, Kristensen K, van Benthem KJ, Magnusson A and others (2017) glmmTMB balances speed and flexibility among packages for zero-inflated generalized linear mixed modeling. The R Journal 9:378–400

Brown CH (1952) Some structural proteins of Mytilus edulis. Journal of Cell Science s3-93:487–502

Brown RL (1967) The use of the embryonic stages of the bay mussel, Mytilus edulis Linnaeus, as a bioassay tool with special reference to sodium pentachlorophenate. Master’s Thesis, Oregon State University

Bulnheim HP, Gosling E (1988) Population genetic structure of mussels from the Baltic Sea. Helgolander Meeresunters 42:113–129

Carrington E (2002) Seasonal variation in the attachment strength of blue mussels: Causes and consequences. Limnology and Oceanography 47:1723–1733

Carthey AJR, Banks PB (2014) Naïveté in novel ecological interactions: Lessons from theory and experimental evidence. Biological Reviews 89:932–949

Carthey AJR, Blumstein DT (2018) Predicting predator recognition in a changing world. Trends in Ecology & Evolution 33:106–115

Cheung SG, Y. TP, M. YK, Shin PKS (2004) Chemical cues from predators and damaged conspecifics affect byssus production in the green-lipped mussel Perna viridis. Marine and Freshwater Behaviour and Physiology 37:127–135

Côté IM (1995) Effects of predatory crab effluent on byssus production in mussels. Journal of Experimental Marine Biology and Ecology 188:233–241

Côté IM, Jelnikar E (1999) Predator-induced clumping behaviour in mussels (Mytilus edulis Linnaeus). Journal of Experimental Marine Biology and Ecology 235:201–211

Crawford BA, Hickman CR, Luhring TM (2012) Testing the threat-sensitive hypothesis with predator familiarity and dietary specificity. Ethology 118:41–48

deRivera CE, Ruiz GM, Hines AH, Jivoff P (2005) Biotic resistance to invasion: Native predator limits abundance and distribution of an introduced crab. Ecology 86:3364–3376

Dzierżyńska-Białończyk A, Jermacz ł, Zielska J, Kobak J (2019) What scares a mussel? Changes in valve movement pattern as an immediate response of a byssate bivalve to biotic factors. Hydrobiologia 841:65–77

Eastwood MM, Donahue MJ, Fowler AE (2007) Reconstructing past biological invasions: Niche shifts in response to invasive predators and competitors. Biological Invasions 9:397–407

Freeman AS, Meszaros J, Byers JE (2009) Poor phenotypic integration of blue mussel inducible defenses in environments with multiple predators. Oikos 118:758–766

Garner YL, Litvaitis MK (2013a) Effects of injured conspecifics and predators on byssogenesis, attachment strength and movement in the blue mussel, Mytilus edulis. Journal of Experimental Marine Biology and Ecology 448:136–140

Garner YL, Litvaitis MK (2013b) Effects of wave exposure, temperature and epibiont fouling on byssal thread production and growth in the blue mussel, Mytilus edulis, in the Gulf of Maine. Journal of Experimental Marine Biology and Ecology 446:52–56

Glude JB (1955) The effects of temperature and predators on the abundance of the soft-shell clam, Mya arenaria, in New England. Transactions of the American Fisheries Society 84:13–26

Gosling E (2003) Bivalve Molluscs: Biology, Ecology and Culture, Vol. Blackwell Publishing Ltd., Oxford, UK

Harley CDG, Randall Hughes A, Hultgren KM, Miner BG, Sorte CJB, Thornber CS, Rodriguez LF, Tomanek L, Williams SL (2006) The impacts of climate change in coastal marine systems. Ecology Letters 9:228–241.

Hatcher A, Grant J, Schofield B (1997) Seasonal changes in the metabolism of cultured mussels (Mytilus edulis l.) from a Nova Scotian inlet: The effects of winter ice cover and nutritive stress. Journal of Experimental Marine Biology and Ecology 217:63–78

Hemmi JM, Merkle T (2009) High stimulus specificity characterizes anti-predator habituation under natural conditions. Proceedings of the Royal Society B: Biological Sciences 276:4381–4388

Hennebicq R, Fabra G, Pellerin C, Marcotte I, Myrand B, Tremblay R (2013) The effect of spawning of cultured mussels (Mytilus edulis) on mechanical properties, chemical and biochemical composition of byssal threads. Aquaculture 410–411:11–17.

Ishida S, Iwasaki K (2003) Reduced byssal thread production and movement by the intertidal mussel Hormomya mutabilis in response to effluent from predators. Journal of Ethology 21:117–122

Jacobsen HP, Stabell OB (1999) Predator-induced alarm responses in the common periwinkle, Littorina littorea: A dependence on season, light conditions, and chemical labelling of predators. Marine Biology 134:551–557

Johnson DS (2015) The savory swimmer swims north: A northern range extension of the blue crab Callinectes sapidus? Journal of Crustacean Biology 35:105–110

Lachance A-A, Hennebicq R, Myrand B, Sévigny J-M and others (2011) Biochemical and genetic characteristics of suspension-cultured mussels (Mytilus edulis) in relation to byssal thread production and losses by fall-off. Aquatic Living Resources 24:283–293

Large SI, Torres P, Smee DL (2012) Behavior and morphology of Nucella lapillus influenced by predator type and predator diet. Aquatic Biology 16:189–196

Le Roux PJ, Branch GM, Joska MAP (1990) On the distribution, diet and possible impact of the invasive European shore crab Carcinus maenas (L.) along the South African coast. South African Journal of Marine Science 9:85–93

Lee CY, Lim SSL, Owen MD (1990) The rate and strength of byssal reattachment by blue mussels (Mytilus edulis L.). Canadian Journal of Zoology 68:2005–2009

Leonard GH, Bertness MD, Yund PO (1999) Crab predation, waterborne cues, and inducible defenses in the blue mussel, Mytilus edulis. Ecology 80:1–14

Lowen JB, Innes DJ, Thompson RJ (2013) Predator-induced defenses differ between sympatric Mytilus edulis and M. trossulus. Marine Ecology Progress Series 475:135–143

Lubchenco J (1978) Plant species diversity in a marine intertidal community: Importance of herbivore food preference and algal competitive abilities. Am Nat 112:23–39

Lurman GJ, Hilton Z, Ragg NLC (2013) Energetics of byssus attachment and feeding in the green-lipped mussel Perna canaliculus. The Biological Bulletin 224:79–88

Matheson K, McKenzie CH, Gregory RS, Robichaud DA, Bradbury IR, Snelgrove PVR, Rose GA (2016) Linking eelgrass decline and impacts on associated fish communities to European green crab Carcinus maenas invasion. Marine Ecology Progress Series 548:31–45

Matoo OB, Lannig G, Bock C, Sokolova IM (2021) Temperature but not ocean acidification affects energy metabolism and enzyme activities in the blue mussel, Mytilus edulis. Ecology and Evolution 11:3366–3379

McBride RS, Tweedie MK, Oliveira K (2018) Reproduction, first-year growth, and expansion of spawning and nursery grounds of black sea bass (Centropristis striata) into a warming Gulf of Maine. Fishery Bulletin 116:323–336

Menge BA (1976) Organization of the New England rocky intertidal community: Role of predation, competition, and environmental heterogeneity. Ecological Monographs 46:355–393

Meyer-Rust KA, Strickland A, Lee B-Y, Sevigny JL, Bradt G, Brown BL (2024) Diet of the blue crab (Callinectes sapidus) during range expansion in Great Bay Estuary, New Hampshire. BMC Genomics 25:1238

Millikin MR, Williams AB (1984) Synopsis of biological data on the blue crab, Callinectes sapidus Rathbun. NOAA Technical Report NMFS 1: FAO Fisheries Synopsis No. 138, US Dept of Commerce, NOAA, NMFS, Washington, DC

Mredul MMH, Sokolov EP, Kong H, Sokolova IM (2024) Spawning acts as a metabolic stressor enhanced by hypoxia and independent of sex in a broadcast marine spawner. Science of The Total Environment 909:168419.

Newcomb LA, Cannistra AF, Carrington E (2022) Divergent effects of ocean warming on byssal attachment in two congener mussel species. Integrative and Comparative Biology 62:700–710

Nye JA, Joyce TM, Kwon Y-O, Link JS (2011) Silver hake tracks changes in Northwest Atlantic circulation. Nature Communications 2:412

Paine RT (1976) Size-limited predation: an observational and experimental approach with the Mytilus-Pisaster interaction. Ecology 57:858–873

Parmesan C, Yohe G (2003) A globally coherent fingerprint of climate change impacts across natural systems. Nature 421:37–42

Pershing AJ, Alexander MA, Brady DC, Brickman D and others (2021) Climate impacts on the Gulf of Maine ecosystem: A review of observed and expected changes in 2050 from rising temperatures. Elementa: Science of the Anthropocene 9:1–18

Petraitis PS (1987) Immobilization of the predatory gastropod, Nucella lapillus, by its prey, Mytilus edulis. The Biological Bulletin 172:307–314

Pinsky ML, Worm B, Fogarty MJ, Sarmiento JL, Levin SA (2013) Marine taxa track local climate velocities. Science 341:1239–1242

R Core Team (2025) R: A language and environment for statistical computing. In: R Foundation for Statistical Computing (ed). R Foundation for Statistical Computing, Vienna, AT

Reimer O, Tedengren M (1996) Phenotypical improvement of morphological defences in the mussel Mytilus edulis induced by exposure to the predator Asterias rubens. Oikos 75:383–390

Reimer O, Tedengren M (1997) Predator-induced changes in byssal attachment, aggregation and migration in the blue mussel, Mytilus edulis. Marine and Freshwater Behaviour and Physiology 30:251–266

Reimer O (1999) Increased gonad ratio in the blue mussel, Mytilus edulis, exposed to starfish predators. Aquatic Ecology 33:185–192

Reimer O, Harms-Ringdahl S (2001) Predator-inducible changes in blue mussels from the predator-free Baltic Sea. Marine Biology 139:959–965

Rickaby R, Sinclair J (2018) Native versus invasive crab effluent effects on byssal thread production in the mussel, Mytilus trossulus (Gould, 1950). The Arbutus Review 9:20–31

Roberts EA, Newcomb LA, McCartha MM, Harrington KJ, LaFramboise SA, Carrington E, Sebens KP (2021) Resource allocation to a structural biomaterial: Induced production of byssal threads decreases growth of a marine mussel. Functional Ecology 35:1222–1239

Roberts EA, Carrington E (2023) Energetic scope limits growth but not byssal thread production of two mytilid mussels. Journal of Experimental Marine Biology and Ecology 567:151927

Ropes JW (1968) The feeding habits of the green crab, Carcinus maenas (L.). Fishery Bulletin 67:183–203

Scattergood LW (1952) The distribution of the green crab, Carcinides maenas (L.) in the north-western Atlantic. Fisheries Circular No. 8, Department of Sea and Shore Fisheries, Augusta, ME

Saurel C, Ng C, Barreau P, Connellan I, Hannon C, Hughes A, Nielsen P (2022) Hatchery protocols for production of blue mussel seeds. AquaVitae.

Sebens KP, Sarà G, Carrington E (2018) Estimation of fitness from energetics and life-history data: An example using mussels. Ecology and Evolution 8:5279–5290

Seed R (1980) Predator-prey relationships between the mud crab Panopeus herbstii, the blue crab, Callinectes sapidus and the Atlantic ribbed mussel Geukensia (=Modiolus) demissa. Estuarine and Coastal Marine Science 11:445–458

Seitz RD, Knick KE, Westphal M (2011) Diet selectivity of juvenile blue crabs (Callinectes sapidus) in Chesapeake Bay. Integrative and Comparative Biology 51:598–607

Sherker ZT, Ellrich JA, Scrosati RA (2017) Predator-induced shell plasticity in mussels hinders predation by drilling snails. Marine Ecology Progress Series 573:167–175

Sih A, Bolnick DI, Luttbeg B, Orrock JL and others (2010) Predator-prey naïveté, antipredator behavior, and the ecology of predator invasions. Oikos 119:610–621

Smith LD, Jennings JA (2000) Induced defensive responses by the bivalve Mytilus edulis to predators with different attack modes. Marine Biology 136:461–469

Sorte CJB, Williams SL, Carlton JT (2010) Marine range shifts and species introductions: Comparative spread rates and community impacts. Global Ecol Biogeogr 19:303–316

Steinback SR, Allen RB, Thunberg E (2008) The benefits of rationalization: The case of the American lobster fishery. Marine Resource Economics 23:37–63

Tan EBP, Beal BF (2015) Interactions between the invasive European green crab, Carcinus maenas (L.), and juveniles of the soft-shell clam, Mya arenaria L., in eastern Maine, USA. Journal of Experimental Marine Biology and Ecology 462:62–73

Venables WN, Ripley BD (2002) Modern applied statistics with S, Vol. Springer, New York, NY, USA

Waite JH (1992) The formation of mussel byssus: Anatomy of a natural manufacturing process. In: Case ST (ed) Structure, Cellular Synthesis and Assembly of Biopolymers. Springer Berlin Heidelberg, Berlin, Heidelberg, p 27–54

Walther G-R, Post E, Convey P, Menzel A and others (2002) Ecological responses to recent climate change. Nature 416:389–395

Young GA (1985) Byssus-thread formation by the mussel Mytilus edulis: Effects of environmental factors. Marine Ecology Progress Series 24:261–271

