## Supplemental Figure for "Non-native crustacean predators conditionally trigger inducible defense behaviors in blue mussels"

**Supplement**

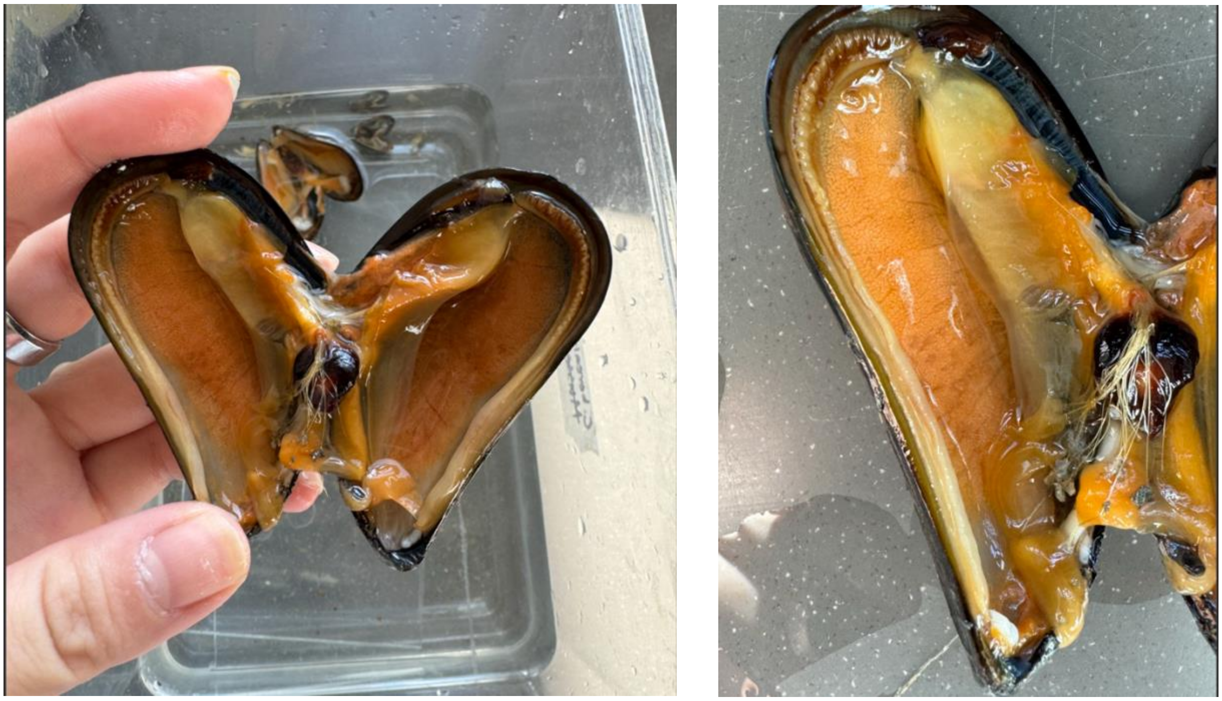

**Supplement 1.** Mytilus edulis female that spawned during trial 11 on November 15, 2024. The shell length of the mussel was 71 mm long.

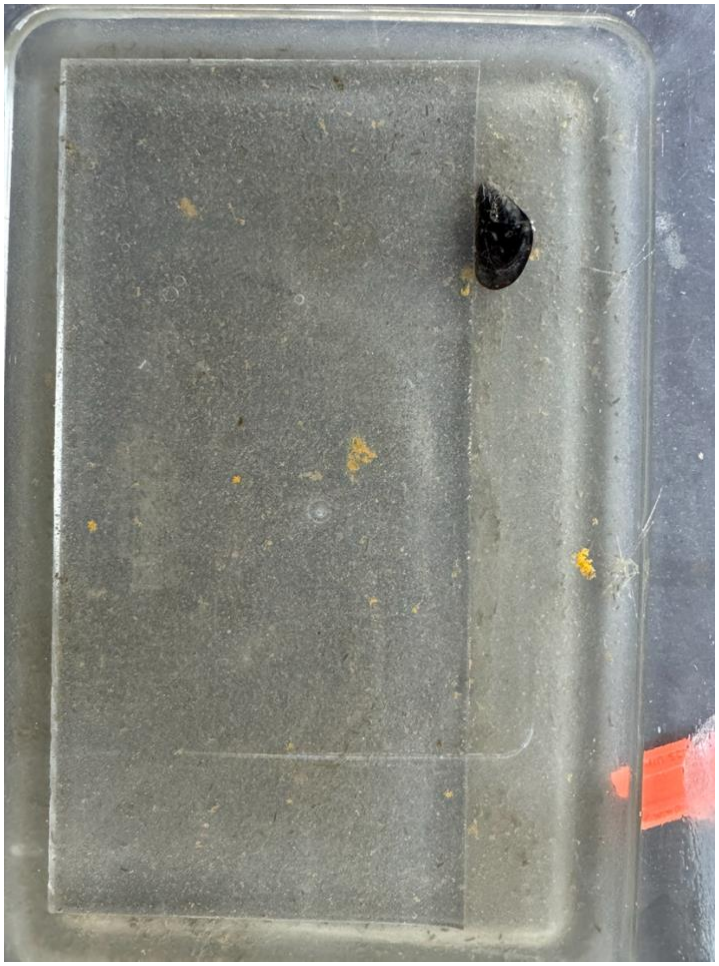

**Supplement 2.** Mytilus edulis female that spawned during trial 10 on November 13, 2024. The shell length of the mussel was 30 mm long. The spawned eggs are aggregated in yellow clusters.
